# Association of cerebellar inflammation and neurodegeneration in a novel spinocerebellar ataxia type 13 mouse model

**DOI:** 10.1101/2024.10.28.620701

**Authors:** Junxiang Yin, Jennifer White, Swati Khare, Michael Wu, Aamir R. Zuberi, Ming Gao, Jerelyn A. Nick, Cathleen M. Lutz, Kyle D. Allen, Harry S. Nick, Michael F. Waters

**Affiliations:** Department of Neurology, Icahn School of Medicine at Mount Sinai, New York,10029, USA; Departments of Neurology, Barrow Neurological Institute, St. Joseph’s Hospital and Medical Center, Dignity Health, Phoenix, Arizona 85013, USA; Department of Neuroscience, University of Florida, Florida 32611, USA; Oxford PharmaGenesis Inc, Newtown, Pennsylvania 18940, USA; Technology Evaluation and Development, The Jackson Laboratory, Bar Harbor, Maine 04609, USA; Rare Disease Translational Center, The Jackson Laboratory, Bar Harbor, Maine 04609, USA; Department of Molecular Genetics and Microbiology, University of Florida, Florida 32611, USA

**Keywords:** neuroinflammation, neurodegeneration, Spinocerebellar ataxia 13, R424H, motor function, cerebellum

## Abstract

**Background:** Neuroinflammation is a recognized pathological characteristic of neurodegenerative diseases. Spinocerebellar ataxia 13 (SCA13) is a progressive neurodegenerative disease with no effective treatments. Our previous studies reported human mutations in *KCNC3* gene are causative for SCA13. Human R423H allelic mutation induces early-onset neurodegeneration and aberrant intracellular retention of Epidermal Growth Factor Receptor (EGFR) in drosophila. However, the neurodegeneration and inflammatory response induced by the R424H allele are unknown in a mammalian model of disease.

**Method:** In this study, a single *Kcnc3* R424H mutation (Analogous to the human SCA13 R423H isoform) transgenic mice were created using CRISPR/Cas 9 technique. Motor function (gait, tremor, coordination and balance) and cerebellar volume (scanned and imaged with 7T MRI) of the R424H transgenic mice were evaluated at multiple timepoints. Neurodegeneration (Purkinje cells loss) as well as cerebellar (astroglia, microglia and macrophage activation) and peripheral (plasma cytokines levels) inflammatory responses were examined and analyzed.

**Result:** The R424H transgenic mice showed marked neurological motor dysfunction with high-frequency tremor, aberrant gait, and short latency to fall in Rotarod testing at 3 and 6 months of age. Abnormal spontaneous firing was recorded in electrophysiology of Purkinje cells. Pathological changes in our R424H transgenic mice included progressive Purkinje cell degeneration and cerebellar atrophy. Over-active microglia, astrocytes, and macrophages were observed in the cerebella of transgenic mice. Pearson correlation analyses indicated that the number of Calbindin positive cells, a Purkinje cell marker, showed a strong inverse correlation with the positive cell number of EGFR, phosphorylated EGFR (pEGFR), and CD68. The expression of EGFR/pEGFR was positively correlated with CD68 and Glial Fibrillary Acidic Protein.

**Conclusion:** Transgenic R424H mice provide a novel SCA13 model showing significant motor deficits, Purkinje cells loss, cerebellar inflammation, and atrophy. Our study suggests that the activation of inflammatory immune cells (astroglia, microglia and macrophages) and strong expression of EGFR/ pEGFR signal in these immune cells are associated with Purkinje cell loss in the cerebellum. This abnormal neuroinflammation may play a significant role in the aggressive procession of neurodegeneration.

## Introduction

Neurodegenerative diseases have gained attention due to the considerable socioeconomic burden they impose. Neuronal loss is a pathological hallmark of several neurodegenerative diseases including Alzheimer’s and Parkinson’s diseases, amyotrophic lateral sclerosis, and spinocerebellar ataxia [1]. It is well documented that inflammatory response plays a critical role in these neurodegenerative diseases [2–6]. Inflammation is not only the consequence of neurodegeneration but also exacerbates the pathological process [7]. Aberrant protein aggregation can induce neuroinflammation, which sequentially aggravates protein aggregation and neurodegeneration[4, 6, 8]. Inflammation has also been observed prior to protein aggregation. In susceptible populations, genetic variations in central nervous system (CNS) cells can induce neuroinflammation [6]. Multiple CNS cells and numerous signaling pathways are involved in the pathogenesis of neurodegeneration [9–12]. Specifically, glial activation has been identified as a feature of neurodegenerative diseases, however, the underlying mechanisms driving neurodegeneration remain largely unclear [4].

Spinocerebellar ataxias (SCA) are a group of neurodegenerative diseases where patients show progressive aberrant gait, tremor, and cerebellar atrophy. There are no effective treatments to slow disease progression [13, 14]. In addition to the progressive neuronal loss, there is increasing evidence that endothelial cells and glia contribute to the pathology of SCAs [15–18]. Inflammatory response in the peripheral and central nervous systems may be linked to the severity of the clinical phenotypes in Spinocerebellar ataxia type 1 (SCA1) [11, 19, 20], 2 [21], 3 [22–26], 6 [27], and 17 [28]. Studies have shown that reactive Bergmann glia may impair neuronal function by interfering with their normal supportive role or triggering neuroinflammation [11]. The distinct subpopulations of cerebellar microglia and oligodendrocytes remain largely unexplored [29, 30]. Previous studies in a SCA1 mouse model have shown that early activation of microglia and astrocytes were closely related with the onset and severity of SCA1[20]. The investigation of neuroinflammation induced by glial activation is therefore important for understanding the mechanism of neurodegeneration in SCAs.

Our previous research indicated that human mutations (R423H, R420H and F448L) in the *KCNC3* gene encoding for the Kv3.3 voltage- dependent potassium channel are causative of SCA13 [31–33]. Clinical characteristics of patients with SCA13 *KCNC3*-R423H are infantile onset, non-progressive cerebellar hypoplasia, lifetime improvement of moto and cognitive function, bradyphrenia, dysarthria, tremor, along with the classic SCA feature of limb, truncal, and gait ataxia [32]. MRI data showed significant cerebellar atrophy in 10-month-old patients with SCA13 pArg423His (human R423H, mouse R424H) [32]. Expression of the human R423H allele in Drosophila markedly induced early-onset neurodegeneration and yielded maldeveloped eyes; expression of the R423H allele in mammalian cells revealed aberrant intracellular retention of epidermal growth factor receptor (EGFR) [32]. Interestingly, EGFR overexpression resulted in a striking rescue of the neurodegeneration and eye phenotype found in this Drosophila model [32]. However, the neurodegenerative mechanism, including the inflammatory response induced by R424H mutation in mice (analogous to the human *KCNC3* R423H isoform), remains unknown.

To understand the molecular mechanism of SCA13, we created a transgenic mouse model with a single R424H mutation using the CRISPR/Cas9 technique. We examined these R424H transgenic mice for motor function and recorded their cerebellar volumes at multiple timepoints. We also evaluated neurodegeneration (defined by Purkinje cell loss) as well as cerebellar (astroglia, microglia, and macrophage activation) and peripheral (plasma cytokines levels) inflammatory response. Results from our analyses confirm that the novel R424H transgenic mouse model shows clinically relevant behaviors and is therefore a robust system to study the pathogenesis of SCA13.

## Materials and methods

### Generation of *Kcnc3* R424H clinical allele using CRISPR/Cas9

To generate the *Kcnc3* R424H mutation transgenic mouse model, we collaborated with the Jackson laboratory to generate the *Kcnc3* R424H clinical allelic mutation through CRISPR/Cas9 targeted FVB/NJ (JR 1800) single cell zygotes. Ensembl *Kcns3*-201 was used as reference transcript for *Kcnc3*. R424 is encoded by CGT and positioned in exon 2. The related sequences were listed in Supplementary Materials. Mouse codon 424 of *Kcnc3* (analogous to the human *KCNC3* R423H isoform) corresponds to human codon 423 of *KCNC3*. Targeting was accomplished by co-microinjection of the S.pyrogenes Cas9 protein with an oligonucleotide RNA guide sequence recognizing the *Kcnc3* antisense DNA strand, 5’-GGGUCAGCUUGAAGAUACGA-3’ and a 132- nucleotide donor DNA fragment containing the three desired nucleotide changes (shown in lower case below) including the R424H missense mutation, corresponding to human R423H; TACTTGGAAGTGGGCCTGTCAGGTCTCAGCTCCAAAGCTGCCAAGGACGTGCTGGGCTTCCTGCGTGT CGTCCGCTTCGTGCGCAT**t**CT**g**C**a**TATCTTCAAGCTGACCCGTCACTTCGTGGGGCTGCGTGTG.

Two other mutations in the genomic sequence were co-introduced. The I422I silent nucleotide change (ATC->ATT) was to eliminate the guide associated AGG PAM sequence to prevent Cas9 recutting (changed from AGG to AGA). The L423L silent nucleotide change (CTT->CTG) was to introduce the TGCA restriction site into the targeted allele, and the R424H mutation change (CGT->CAT) was to introduce the desired missense mutation into the genome. All three nucleotide changes are present in the targeted JR 29698 mouse strain. Strain nomenclature JR 29698 FVB-*Kcnc3^em2Lutzy^*/J. We utilized this forward primer: TGTGGTCTGGTTCACCTTTG and reverse primer: CTTGAAGTAGGTGTGGTTGGAG for genotyping PCR to confirm the presence of the mutation.

All mice were housed in a temperature and humidity-controlled vivarium, kept on a 12-hour dark/light cycle, and had free access to food and water. All experimental procedures were approved by the Institutional Animal Care and Use Committee of the Barrow Neurological Institute and performed according to the Revised Guide for the Care and Use of Laboratory Animals. All principles of laboratory animal care (NIH publication No. 86-23, revised 1985) were followed.

### Tremor analysis

1, 3, and 9 -month-old mice were held in a scruff and handled by the same person for each trial. High speed video recordings (2 trials, 8 seconds per trials, n=5 per group) were taken under ventral view at the frame rate of 240 frames per second, using a Sony Cyber-shot DSC-RX100 IV digital camera. DLT data viewer (https://biomech.web.unc.edu/dltdv) was utilized by a blinded observer for manually marking the position of the left front paw middle digit for a single animal in each frame [34]. The periodogram function of MATLAB was utilized along with in-house code to find the frequency of movement of the paw.

### Spatiotemporal gait analysis

Given the poor performance of R424H mice in the complex Rotarod coordination task, we chose to examine gait abnormalities using a simplified task associated with a lower level of stress. We performed a quantitative spatiotemporal gait analysis of 3-month-old mice walking across an arena with a system involving high-speed videography [35]. This approach increased hardware sensitivity with high-speed videography (250 frames per second) and reduced the contribution of animal stress to gait measurements by avoiding a treadmill and incorporated a second angle of view offering estimation of both spatial and temporal parameters. We manually digitized two-time variables (toe-off and foot-strike events) for each paw, and location coordinates (x,y) for each step. Animals underwent gait testing as described by Dr. Kyle D. Allen’s group [36]. The animals were acclimatized to the testing room for 1 hour and allowed to explore the gait arena without any external stimuli until 3 trials each were collected where the animal walked with an approximate constant velocity. High speed videography was utilized to collect walking videos at 250 frames per second (RedLake, M3, DEL Imaging Systems LLC, NH). The collected videos were hand digitized by an observer using a modified version of DLT data viewer in MATLAB [34]. Digitized data was then processed using MS Excel to calculate velocity, stride length, stance times, step widths along with spatial and temporal gait symmetry using methods described earlier [35]. Velocity was included as a covariate for calculations of stride length, step width and percentage stance time analyses as per previous studies [35], and calculated residuals were utilized for statistical analyses.

### Rotarod test

The rotarod test has been used to assess motor coordination and balance alterations. The rotarod apparatus consists of a rod which rotates at an adjustable speed. The speed of the rod increases with time, and the amount of time the animal remains on the device is recorded. In this study, rotarod data was obtained using a Rotarod (IITC Life Science Rotarod Series 8) for mice at the ages of 3 and 6 months. Mice were acclimated to the procedure room for 1 h, then trained for three consecutive days and tested for one day. Data from four trials per animal were acquired on the fourth day of testing. The test length was set to 10 minutes, and the rod was accelerated from 4 rpm to 40 rpm over 5 minutes. Latency to fall was recorded, and mice were given 15 min of rest between each trial.

### Magnetic Resonance Imaging (MRI) and brain volume measurement

Mice aged 1, 3, and 9 months were scanned by MRI for brain volume measurement. MR images were acquired on a Bruker Biospec 7.0T small animal MR scanner (Bruker Medizintechnik, MA) with a 72mm transmit coil and a surface receive coil. We employed a 3D RARE imaging sequence with RARE factor of 8, 4 averages, TR/TEeff= 500 ms/28 ms; matrix size of 180 × 128 × 88; FOV 16 mm× 12.8 mm× 8.8 mm; yielding 100 μm isotropic voxels with 44 min scan time. A sagittal multiple slice 2D T2- weighted RARE sequence was acquired as a reference image (with RARE factor of 8, 10 averages, TR/TEeff= 5000 ms/60 ms; matrix size of 220 × 100; FOV 22 mm× 10 mm; 25 slices) to visualize sagittal morphology. A gas mixture of oxygen and isoflurane (1–2%) was used for anesthesia maintenance. The respiratory rate and body temperature were monitored constantly with an MR-compatible small animal monitoring system (PC_SAM, model #1025, SA Instruments, NY). The body temperature of the mouse was maintained at 36.5°C using water circulation imbedded into the animal cradle, and the respiratory rate was maintained around 50–65 bpm by adjusting the concentration of the isoflurane gas mixture.

3D MR Bruker files were converted to a NIfTI format in ImageJ v.1.51j8. The converted 3D MR images were skull-stripped using the Brain Surface Extractor within BrainSuite v.17a to remove all non-brain material [37]. Inaccuracies were corrected in all three axes by manually editing the masks using Brain Suite v.17a [37]. After skull stripping, the cerebella were manually extracted from the brains using a routine allowing manual operator contouring. The volumes of the whole brains and cerebella were calculated in Brain Suite v.17a using the region of interest (ROI) volume estimation function.

### Electrophysiology

Sagittal cerebellar slices (300-μm thickness) were prepared from 1-month-old mice (wild-type: 3-4 slices per mouse, n=7, and R424H: 3-4 slices per mouse, n=5). The slices were then incubated for at least 1 hour in a preincubation chamber (Warner Instruments, CT) at room temperature in conventional artificial CSF containing (in mM): 125 NaCl, 3 KCL, 2 CaCl2, 1 MgCl2, 1.25 NaH2PO4, 26 NaHCO3 and 10 glucose, continuously saturated with 95% O2 and 5% CO2. Cell-attached recordings were acquired with a MultiClamp 700B Microelectrode Amplifier in voltage-clamp mode at 33–34°C, by visually identifying Purkinje cell somata in anterior lobule III using Zeiss Axio Imager 2 Microscope. The resistance of the electrode was 4 to 5 MΩ when filled with internal solution (in mM: 140 potassium gluconate, 5 KCl, 10 HEPES, 0.2 EGTA, 2 MgCl2, 4 MgATP, 0.3 Na2GTP and 10 Na2- phosphocreatine, pH 7.3 with KOH). The holding potential was 0 mV. Signals were filtered at 2 kHz and sampled at 10 kHz. On-line data acquisition and off-line data analysis were performed using Clampfit.

### Plasma and brain collection, tissues sectioning

Mice were deeply anesthetized with isoflurane, plasma was isolated from peripheral blood collected via transcardiac puncture, mice were then perfused with cold PBS using a Watson-Marlow 120S pump. Half of the cerebellum was subsequently collected and immersed in 4% paraformaldehyde for fixation at 4⁰C overnight. The cerebella were then subjected to several consecutive 30% sucrose treatments until they sank to the bottom of the containment tube, after which they were embedded separately in the Scigen Tissue-Plus™ OCT. 16µm sections were cut using a Leica CM3050 S Cryostat. Sections were mounted on Superfrost Plus microscope slides and stored at -20°C prior to immunohistochemical staining. The other half of the cerebella was subsequently collected in cryoprotectant tubes, then flash frozen in liquid nitrogen, and stored at -80°C for western blot analysis.

### Immunofluorescence staining and imaging

Cryostat sections were thawed in a humid chamber at room temperature for 30 minutes and washed three times for 10 minutes each in PBS (phosphate buffered saline) with tween 20 0.1% (PBST). The sections were blocked with 5% normal goat serum (NGS) in PBS triton X-100 0.1% at room temperature for 1 hour. They were then subjected to the following primary antibodies: anti-EGFR (Cell signaling, #4267, 1:100), anti-phospho-(Tyr1068)-EGFR (Cell signaling, #2234, 1:100), anti-CD68 (Cell signaling, #97778, 1:100), and anti-cleaved caspase 3 (CC3, Cell signaling, #9664, 1:100). Sections were incubated overnight at 4⁰C followed by three 10 min PBST washes. Next, they were incubated in secondary antibody (Alexa Fluor 555, Invitrogen, 1:1000) for 1 hour at room temperature and washed again 3 times in PBST. In the last step slides were incubated overnight the following conjugated antibodies: anti-Calbindin 488 (Cell signaling, #13176, 1:100), anti-Glial Fibrillary Acidic Protein (GFAP) Alexa Fluor 488 (Invitrogen, A-21294, 1:200), Ionized calcium binding adaptor molecule 1 (Iba1) Alexa Fluor 488 (Cell Signaling, #20825, 1:100), and CD68 Alexa Fluor 488 (Cell Signaling, #51644, 1:200). Slides were counterstained with DAPI (Thermo Scientific, 1:1000). Slides were washed twice in PBS for 20 minutes each, with one last wash in PBS for 20 minutes. Slides were sealed with coverslips using VECTASHIELD Antifade Mounting Medium (H-1000m, Vector Laboratories,CA) and the edges were sealed using a clear quick dry nail polish. Slides were imaged using a Nikon A1R HD25 Confocal. Cell count and mean intensities were collected using the NIS-Elements AR software and analyzed in GraphPad Prism. For cell counts, three 200x200 pixel Region of interests (ROIs) were drawn and placed over three representative areas within a 10x image taken from the cerebellum of each animal. Cells were hand counted within each ROI and the average between all three ROIs was calculated for further analysis. Mean intensities were collected by making a rolling ball correction for each cerebellar image to correct for background, then reading the intensity for the channel of interest calculated by the NIS-Elements software.

### Western blot

Brain tissue was homogenized in RIPA buffer (Thermo Scientific, Rockford, IL) with a protease and phosphatase inhibitor cocktail (Sigma) on ice. The protein concentrations were determined using a BCA protein assay kit (Thermo Scientific, Rockford, IL). Western blot analysis was performed with 20 µg (cleaved caspase 3, CD68, pEGFR) and 50 µg total protein (Calbindin and GFAP) extract separated on 8-16% and 4-20% Tris-Glycine gels (Invitrogen, Carlsbad, CA) that were subsequently transferred to nitrocellulose membrane (Millipore, Billerica, MA). Membranes were blocked with 5% nonfat dry milk or bovine serum albumin in TBST and washed in TBST. Membranes were incubated with anti-calbindin (Cell signaling, #65152, 1:1000), anti-phospho-(Tyr1068)-EGFR (Cell signaling, #2234, 1:1000), anti-GFAP (Cell Signaling, 80788, 1:1000), anti-CD68 (Cell signaling, #97778, 1:1000), anti-Cleaved Caspase 3 (CC3, Cell signaling, #9664, 1:1000) at 4°C overnight. They were washed three times for 10 minutes each in PBST at RT, then incubated in secondary antibody (LI-COR, IRDye 800CW goat anti-rabbit IgG or IRDye 680RD goat anti-mouse IgG, 1:10000) for 1 hour at RT.

Normalization of results was ensured by running a control in the form of β-actin (Cell Signaling, #3700, 1:5,000). Membranes were scanned, and the density was quantified on the LI-COR Odyssey XF imaging system. The data were presented as a percentage of target protein relative to β-actin.

### Plasma proinflammatory interleukins and cytokines measurement

Plasma was collected from wild-type mice and R424H mice at age 1, 3, and 6 months. Plasma proinflammatory interleukins and cytokines were then quantitatively measured using Meso Scale Discovery (MSD) V-PLEX Mouse Cytokine 19-Plex Kit (MSD, #K15255D) according to the manufacturer’s exact protocol. Developed under rigorous design control, MSD V-PLEX kit provides accurate and reproducible results. The V-PLEX Mouse Cytokine 19-Plex Kit includes 19 biomarkers associated with the inflammatory response and immune system regulation, these 19 biomarkers include interleukin-1 beta (IL-1β), IL-2, IL-4, IL-5, IL-6, IL-9, IL-10, IL-12p70, IL-15, IL-17A/F, IL-27p28/IL-30, IL-33, Interferon gamma (IFN γ), Interferon gamma-induced protein 10 (IP-10), KC/GRO protein (interleukin-8 related protein), Monocyte chemoattractant protein 1 (MCP-1), Macrophage inflammatory protein 1 alpha (MIP-1α), Macrophage inflammatory protein 2 (MIP-2), Tumor necrosis factor alpha (TNF-α).

## Statistical analysis

All data are presented as mean ± standard error of mean. For spatiotemporal gait analysis, residual changes were calculated for stride length, stance time and step widths with velocity as a covariate and wild-type mice as controls in MS Excel. Resulting spatiotemporal gait data and other behavioral analysis data was compared using the Mann Whitney U-test or unpaired t-test with Welch’s correction in GraphPad Prism. For comparisons involving multiple time-points and genotypes, one-way ANOVA was performed in GraphPad Prism. Statistical significance was defined as p < 0.05 for all analyses.

## Results

### Generation of R424H mice

To generate the Kcnc3 R424H point mutation transgenic model mouse, we utilized CRISPR/Cas9. A total of three mutations in the genomic sequence were introduced, including mouse allele Kcnc3 I422I, L423L, R424H. The changes introduced were ATC/CTT/CGT/ -> ATT/CTG/CAT/. The I422I silent nucleotide change (ATC->ATT) was to eliminate the guide associated AGG PAM sequence to prevent Cas9 recutting (changed from AGG to AGA). The L423L silent nucleotide change (CTT->CTG) was to introduce the TGCA restriction site into the targeted allele, and the R424H mutation change (CGT->CAT) was to introduce the desired missense mutation into the genome (Fig 1.A). The modified allele contains the methylation dependent CviRI restriction enzyme sequence (TGCA) and is present in the *Kcnc3* R424H mutant. Genotyping PCR confirmed there was a different DNA band between wild type and transgenic mice (Fig 1.B).

**Fig 1.**
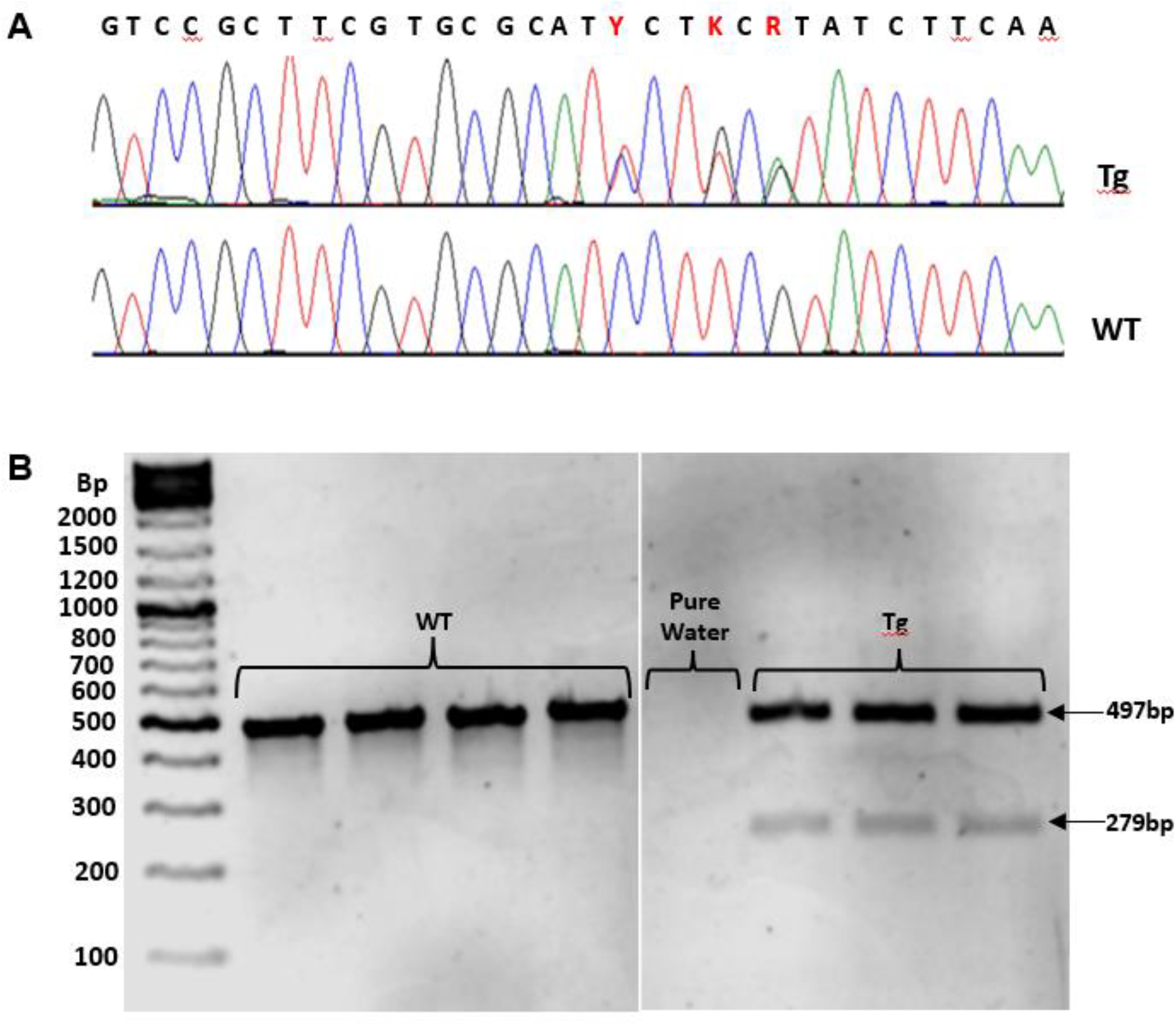
Generation of R424H mice. (A) The genomic sequence recognizing point mutations (red letter) at Kcnc3 I422I (ATC->ATT), L423L (CTT->CTG), R424H (CGT->CAT) from Wild type mouse (WT) and R424H mouse (Tg). (B) Identification of Tg mice and WT mice with using PCR genotyping.

### R424H mutation induced neurological deficits

We evaluated the motor function of R424H mice including spatiotemporal gait pattern, tremor, and Rotarod performance. Single R424H mutation mice displayed aberrant gait and an abnormal power spike in frequency of tremor as early as one month of age. 3-month-old transgenic mice showed high frequency tremor with a powerful spike at 42.69 ± 4Hz, as compared to age-matched wild-type mice that did not exhibit this frequency of limb movement. (Fig 2. A, B). Analysis of spatiotemporal gait patterns indicated that the transgenic mice displayed multiple aberrations. The transgenic mice exhibited a significantly lower velocity and notably different residual stride lengths in comparison with wild-type mice at 3 months (Fig 2. C, D). There were also significant differences in step widths of both fore and hind limbs between mutant mice and wild type (Fig 2. E, F), the recording gait videos were listed in Supplementary Materials. The Rotarod testing was used to assess balance, grip strength and motor coordination of mice. At 3 and 6 months, the transgenic mice exhibited significantly shorter latency to fall compared to those of wild-type mice (Fig 2. G, H).

**Fig 2.**
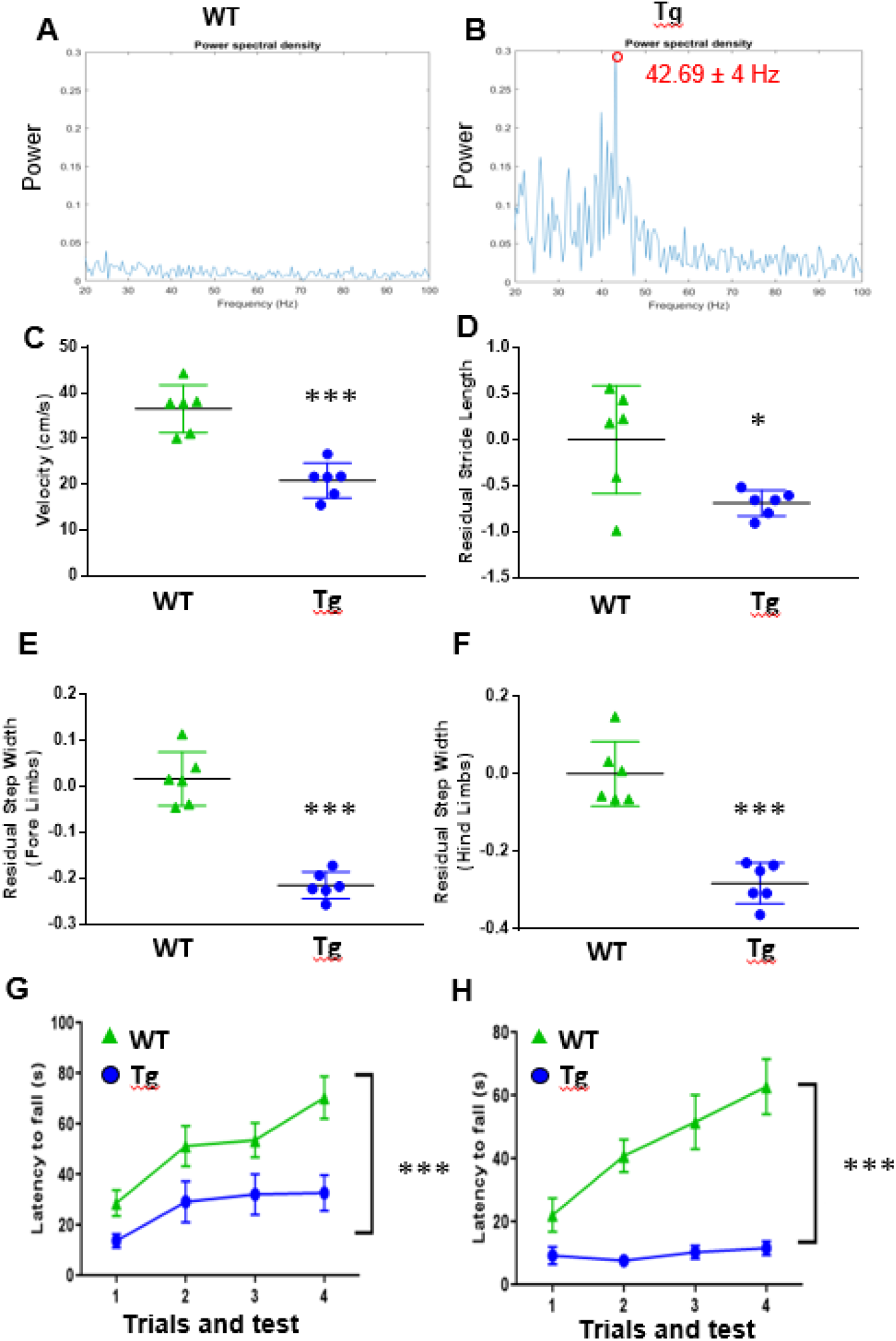
R424H mutation induced neurological deficit. Motor function of R424H (Tg) mice including spatiotemporal gait pattern, tremor, and Rotarod performance was evaluated at different age time points.(A-B) Tremor analysis in wild-type (WT) control mice and Tg mice: (A-B) Periodograms describe the relative power of frequencies present in data collected from fore paw tremors of wild-type (n=6, trials = 2) and Tg mice (n=6, trials = 2). The powerful spike of frequencies is difference between the wild-type mice (A, normal power spectrum), and Tg mice (B, an abnormal power spike in frequency at 42.69±4Hz). (C-F) Spatiotemporal gait analysis in WT mice and Tg mice: (C) Velocities of locomotion, (D) Residual stride length (cm), (E) Residual step widths (cm) of fore, and (F) hind limbs. Rotarod testing in WT mice and Tg mice: (G) Rotarod was evaluated for mice at ages of 3 months (wild-type mice n = 14, Tg mice n=11) and (H) 6 months (WT mice n = 15, Tg mice n=9). Note: Data was presented as mean ± SEM and unpaired t-test or one-way ANOVA Turkey multiple comparison test was used in GraphPad Prism 10. * p <0.05, *** p <0.001.

### R424H mutation caused progressive cerebellar atrophy

In this study, we used 7T MRI to evaluate the brain volumes in mice at ages of 1, 3 and 9 months (Fig 3 A-F). Our data indicated that the cerebellar volumes of transgenic mice were significantly smaller (p < 0.05 or 0.01), compared to those of wild type mice in all three age groups. Total brain volume was only significantly different at the 9-month age point, indicating that this change was progressive with age. (Fig 3. G, H).

**Fig 3.**
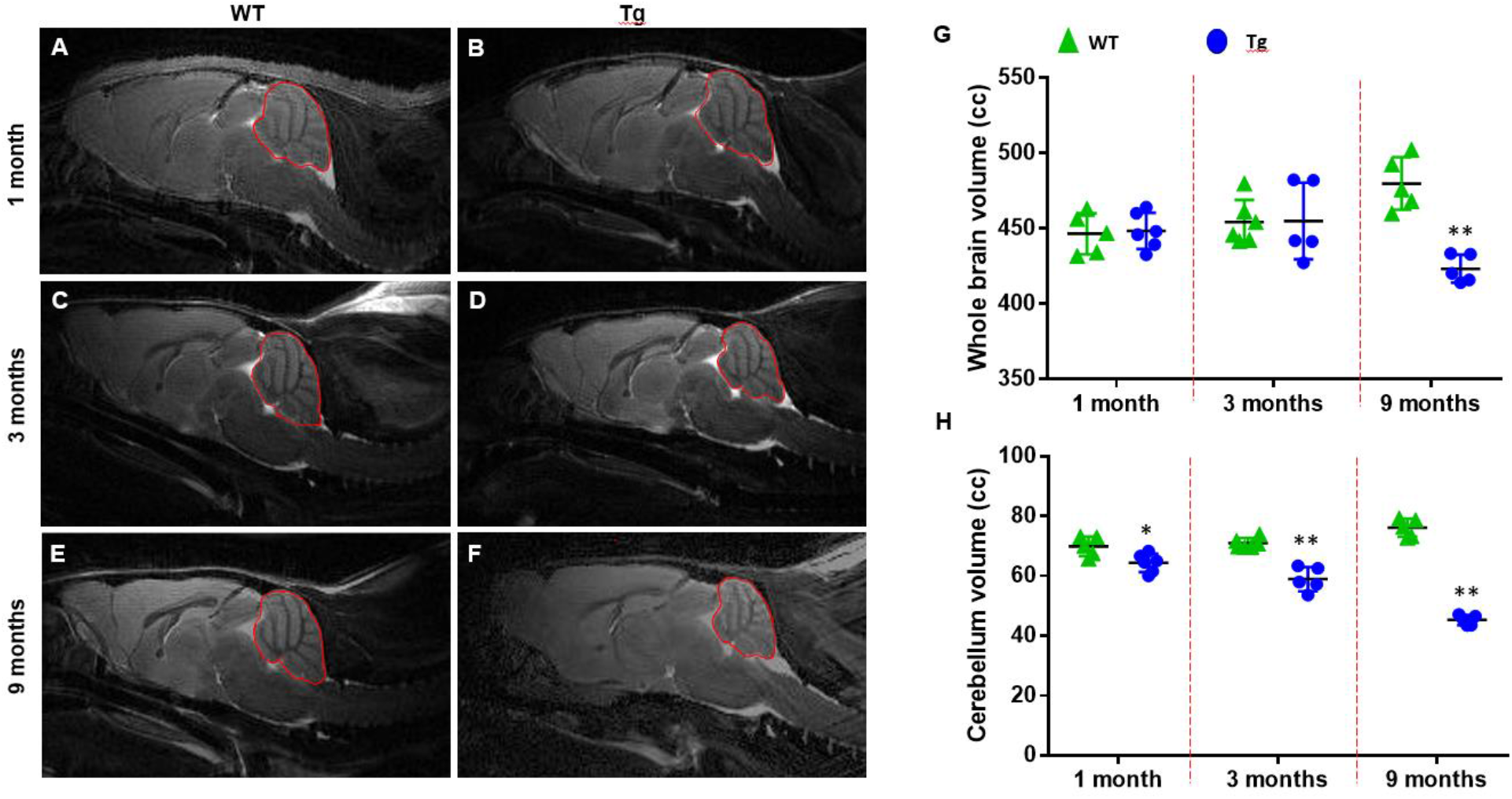
R424H mutation caused progressive cerebellar atrophy. Representative 7T MRI images of wild-type (WT) and R424H (Tg) mice at the ages of 1 month (A, B), 3 months (C, D), and 9 months (E, F). (G) The brain volumes of mice at 1 month (n = 5-6), 3 months (n = 5-6) and 9 months (n = 4). (H) The cerebellar volumes of mice. Different mice were used, and data was collected at every time point. Note: Data was presented as mean ± SEM and analyzed using ordinary one-way ANOVA in GraphPad Prism 10. * p <0.05, ** p <0.01.

### R424H mutation induced abnormal spontaneous firing of Purkinje cells

The loss of *Kcnc3* in Purkinje cells resulted in broadening of action potentials [38, 39]. *Kcnc3* R424H expression through lentiviral transduction revealed altered physiology and pathology of the cerebellar Purkinje cells with a degeneration in the viability in mouse primary cultured Purkinje cells [40]. Electrophysiological recordings of these transduced cells showed severe suppression of current amplitude [40]. In our new R424H transgenic mice, we confirmed that there was aberrant spontaneous firing of Purkinje cells during the electrophysiological recordings in sagittal cerebellar slices at the age of 1 month (Fig 4.). The firing frequency of Purkinje cells in transgenic mice was significantly reduced while the variation of inter-spike interval was significantly increased in comparison with that of the wild-type mice, confirming a change of firing pattern in transgenic mice (Fig 4.).

**Fig 4.**
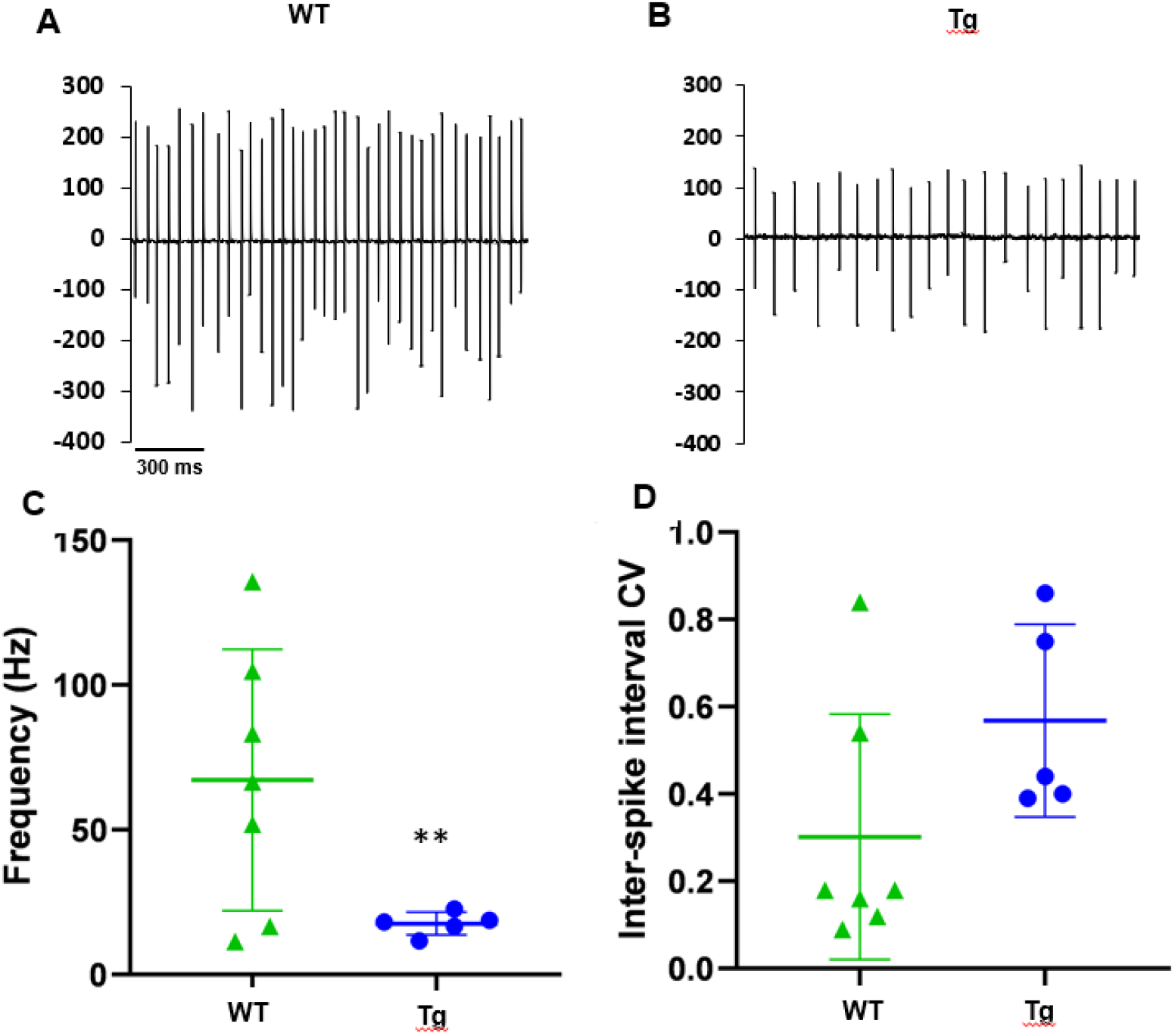
R424H mutation induced abnormal spontaneous firing of Purkinje cells. Sagittal cerebellar slices were prepared from 1-month-old mice. Cell-attached recordings were acquired with a Multiclamp 700B amplifier in voltage-clamp mode by visually identifying Purkinje cell somata in anterior lobule III using an upright microscope (Zeiss). On-line data acquisition and off-line data analysis were performed using Clampfit. (A) Representative trace from wild-type (WT) mice cerebellar slices, (B) Representative trace from R424H (Tg) mice cerebellar slices. (C) Spontaneous firing frequency of Purkinje cells, (D) Co-efficient of variation (CV) of the Inter-spike intervals (ISI) of Purkinje cells. Note: Data was presented as mean ± SEM and analyzed using unpaired t-test in GraphPad Prism 10. WT mice, 4 slices per mouse, n=7, and Tg mice: 4-5 slices per mouse, n=5. ** p <0.01.

### R424H mutation induced cerebellar progressive astrogliosis and apoptosis

Neuroinflammation is associated with the pathogenesis of neurodegenerative diseases. Three major types of glial cells in the central nervous system include astrocytes, microglia, and oligodendrocytes. Hyperactivated cerebellar astrocytes are considered to be one of the earliest responses to the abnormal behavior of neurons induced by ataxia-related gene mutations [41]. In our novel R424H mutation transgenic mouse model, immunofluorescence imaging showed strong GFAP signals in both the molecular layer and white matter of cerebella of mice at ages of 1, 3, and 6 month(s) (Fig 5.A). Additionally, the mean intensity of GFAP positive signal was significantly higher (p<0.01) in cerebella of 3-month-old R424H mice compared to age-matched wild type mice (Fig 5.B, D, E). Our MRI data indicated progressive cerebellar atrophy in the R424H transgenic mice, suggesting that there was neuronal injury and loss. Thus, we investigated whether expression of cleaved caspase 3, an apoptosis marker, was different in the cerebella of the transgenic mice. The immunofluorescence images clearly showed that there was strong cleaved caspase 3 positive signal in the molecular layer and Purkinje cell layer of R424H transgenic mice cerebella at the ages of 3 and 6 months (Fig 5.A). The mean intensity of cleaved caspase 3 positive signal was significantly higher (p<0.01) in the cerebella of R424H mice at 3 and 6 months in comparison with age-matched wild type mice (Fig 5.C, D, E).

**Fig 5.**
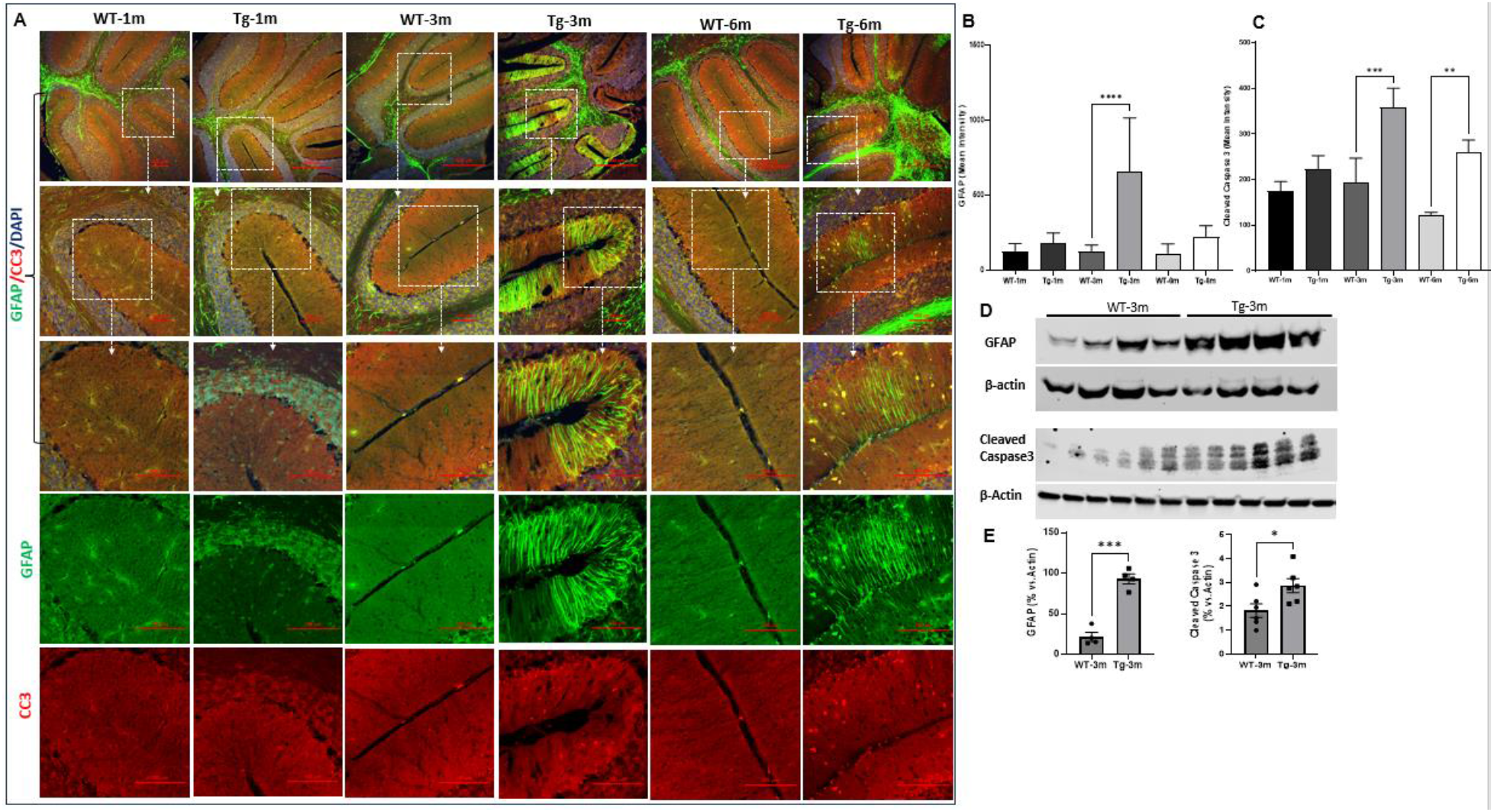
R424H mutation induced cerebellar progressive astrogliosis and apoptosis. To examine astrogliosis and apoptosis in the cerebellum, the location and protein levels of Glial Fibrillary Acidic Protein (GFAP) and Cleaved Caspase 3 (CC3) were examined in the cerebella of mice at different time points. Half of cerebella of mice were fixed for immunofluorescence staining, other half of the cerebella were fresh frozen for western blot analysis. (A) Images representatives of anti-GFAP and anti-CC3 at the ages of 1,3, and 6 months. (B) Quantitative analysis of mean intensity of immunofluorescence staining signal of GFAP. (C) Quantitative analysis of mean intensity of immunofluorescence staining signal of CC3. (D) Western blot images of GFAP and CC3. (E) Quantitative analysis of western blot of GFAP and CC3. Note: Data was presented as mean ± SEM and unpaired t-test or one-way ANOVA Turkey multiple comparison test was used for quantitative analysis in GraphPad Prism 10. * p <0.05, ** p <0.01, *** p <0.001, **** p <0.0001.

### R424H mutation significantly activated microglia and increased macrophage infiltration in the cerebella of transgenic mice

Microglia are the primary immune cells of the CNS and play a significant role in etiology and pathogenesis of neurodegenerative disease. They are the first cells to respond to neuroinflammation or damage in the brain [42]. We evaluated the expression of Iba-1, a microglia marker, in the CNS of our SCA mouse model in comparison to age-matched wild type mice. Our images showed that Iba-1 positive signals were both more frequent and of a stronger intensity in R424H transgenic mice at the ages of 3 and 6 months (Fig 6.A, B). Interestingly, our data also showed there was a strong expression of EGFR in the molecular layer colocalized with Iba-1 signaling from microglia (Fig 6.A, C). This suggests that the aberrant expression of EGFR was associated with activated microglia. The cluster of differentiation 68 (CD68) is a 110 kDa transmembrane glycoprotein that is widely expressed in macrophages and microglia [43]. CD68 plays an essential role in the process of inflammation and auto-immunity [44]. Our images showed there were strong CD68 positive signals in whole cerebella (molecular layer, granular cells layer, and white matter) of R424H transgenic mice at the ages of 3 and 6 months (Fig 7.A). These findings correspond with the obvious and progressive loss of Purkinje cells observed in these transgenic mice at 3 and 6 -month timepoints (Fig 7.A, C, D, E). Our statistical data indicated that the number of CD68 positive cells was significantly increased while the number of Calbindin (a Purkinje cell marker) positive cells markedly declined in transgenic mice at 3 and 6 months compared with that of age-matched wild type mice (Fig 7.E).

**Fig 6.**
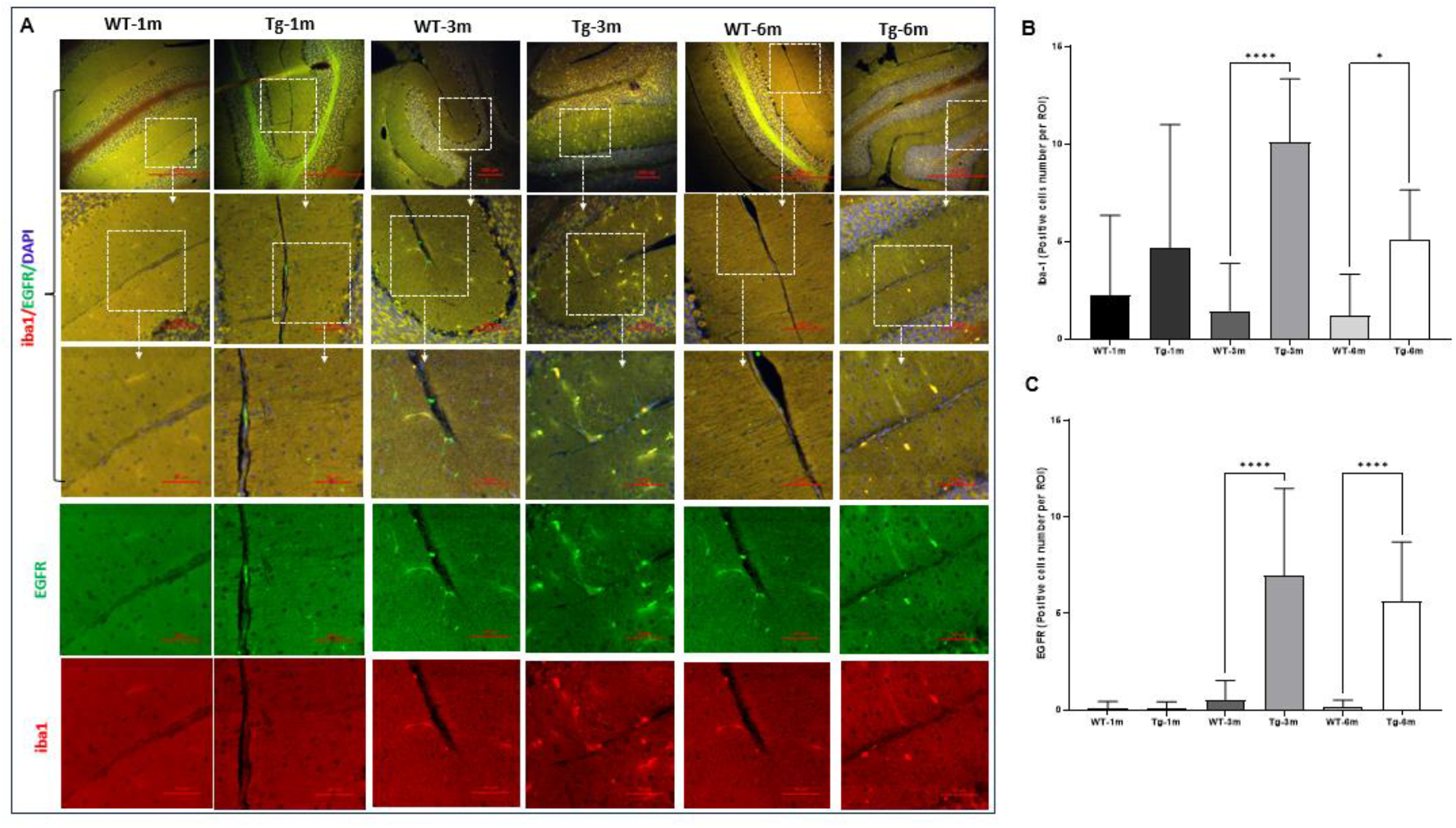
R424H mutation induced cerebellar progressive microglia activation with EGFR colocalized expression. To examine microglia activation and EGFR expression in the cerebellum, the location of Iba1 and EGFR were examined in the cerebella of mice at different time points. (A) Images representatives of anti-Iba1 and anti-EGFR at the ages of 1,3, and 6 months. (B) Quantitative analysis of positive signal cells number of immunofluorescence staining of Iba1. (C) Quantitative analysis of positive signal cells number of immunofluorescence staining of EGFR. Note: Data was presented as mean ± SEM and one-way ANOVA Turkey multiple comparison test was used for quantitative analysis in GraphPad Prism 10. * p <0.05, **** p <0.0001.

**Fig 7.**
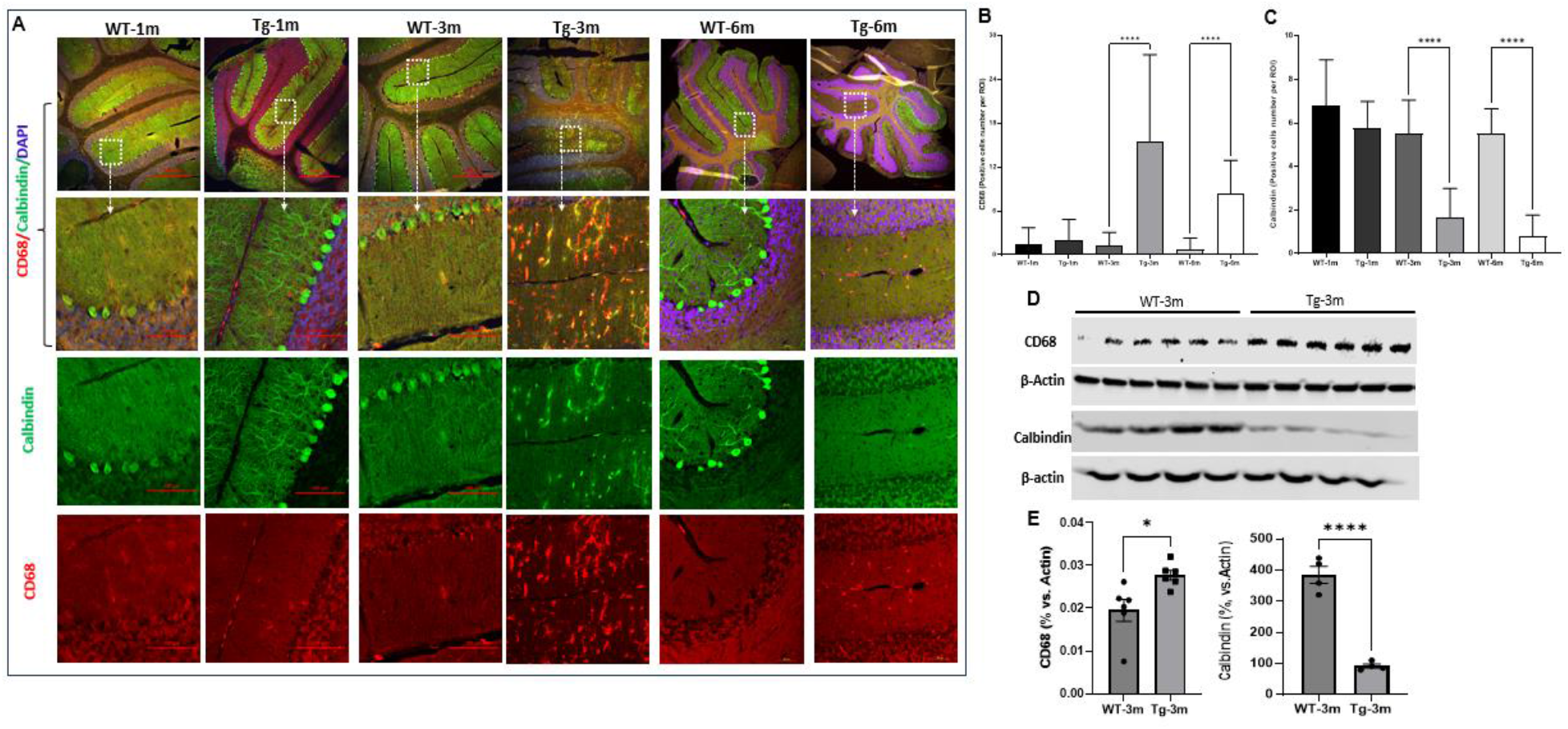
R424H mutation activated cerebellar macrophage (CD68+ cells) and induced Purkinje cells loss. To examine macrophage activation and Purkinje cells loss in the cerebellum, the location and protein levels of CD 68 (macrophage marker) and Calbindin (Purkinje cell marker) were examined in the cerebella of mice at different time points. Half of cerebella of mice were fixed for immunofluorescence staining, other half of the cerebella were fresh frozen for western blot analysis. (A) Images representatives of anti-Calbindin and anti-CD68 at the ages of 1,3, and 6 months. (B) Quantitative analysis of positive cells number of immunofluorescence staining signal of CD68. (C) Quantitative analysis of positive cells number of immunofluorescence staining signal of Calbindin. (D) Western blot images of CD68 and Calbindin. (E) Quantitative analysis of western blot of CD68 and Calbindin. Note: Data was presented as mean ± SEM and unpaired t-test or one-way ANOVA Turkey multiple comparison test was used for quantitative analysis in GraphPad Prism 10. * p <0.05, **** p <0.0001.

### R424H mutation induced aberrant EGFR/pEGFR expression in cerebellar molecular layer CD68+ cells

Our previous studies showed that the R424H mutation in drosophila markedly induced early-onset neurodegeneration and aberrant intracellular retention of epidermal growth factor receptor (EGFR). Interestingly, EGFR overexpression resulted in a striking rescue of the neurodegeneration and eye phenotype found in this drosophila model [32]. EGFR belongs to the receptor tyrosine kinase superfamily [45–47] and it plays a fundamental role during various cellular responses throughout the central nervous system [45, 46, 48]. Historically, the role of EGFR in the nervous system has been underestimated and poorly investigated [48]. Recent studies showed that hyperphosphorylation of EGFR resulted in its neurotoxic intracellular aggregation [49–53]. Targeting EGFR activation showed therapeutic benefits in neurodegenerative diseases and brain injuries [54–58]. In this study, we evaluated the protein expression of EGFR and phosphorylated (Tyr1068) EGFR in the cerebella of mice. We found that EGFR and pEGFR were strongly expressed in the cerebella of transgenic mice at the ages of 3 and 6 months (Fig 6, Fig 8, and Fig 9.). Additionally, EGFR and pEGFR immunoreactivities were colocalized with Iba-1 or CD68 positive microglia and macrophages, specifically in the molecular layer of cerebella. Quantitative evaluations showed that the number of EGFR and pEGFR positive cells were notably increased in transgenic mice at 3 and 6 months compared with that of age-matched wild type mice (Fig 6.C and Fig 9. B). These data suggested that EGFR/pEGFR signaling may play an important role in microglia and macrophage activation.

**Fig 8.**
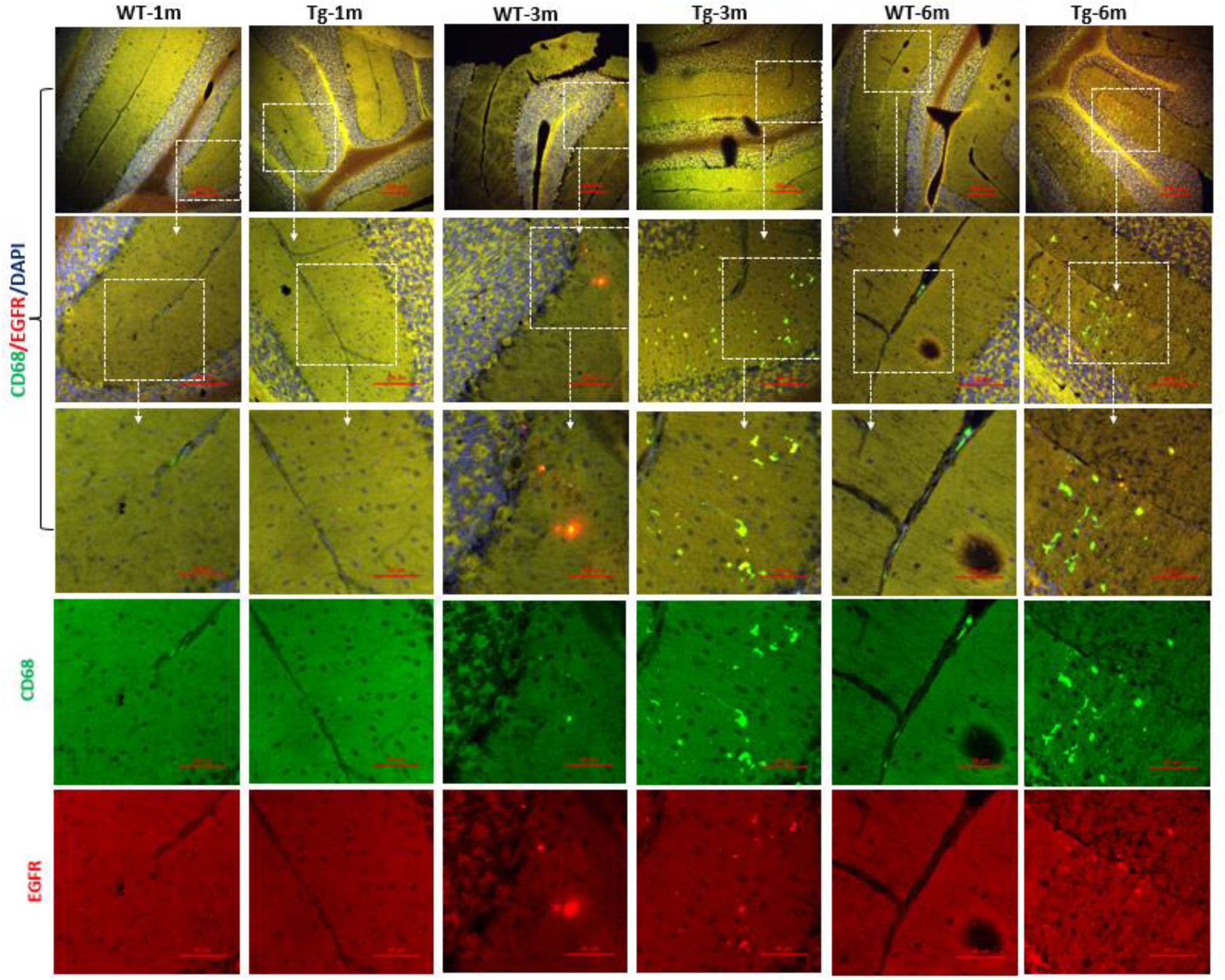
R424H mutation induced cerebellar EGFR overexpression in macrophage. To examine EGFR expression in macrophage, the colocalization of CD68 and EGFR were investigated in the cerebella of mice at different time points. Images representatives of anti-CD68 and anti-EGFR at the ages of 1,3, and 6 months.

**Fig 9.**
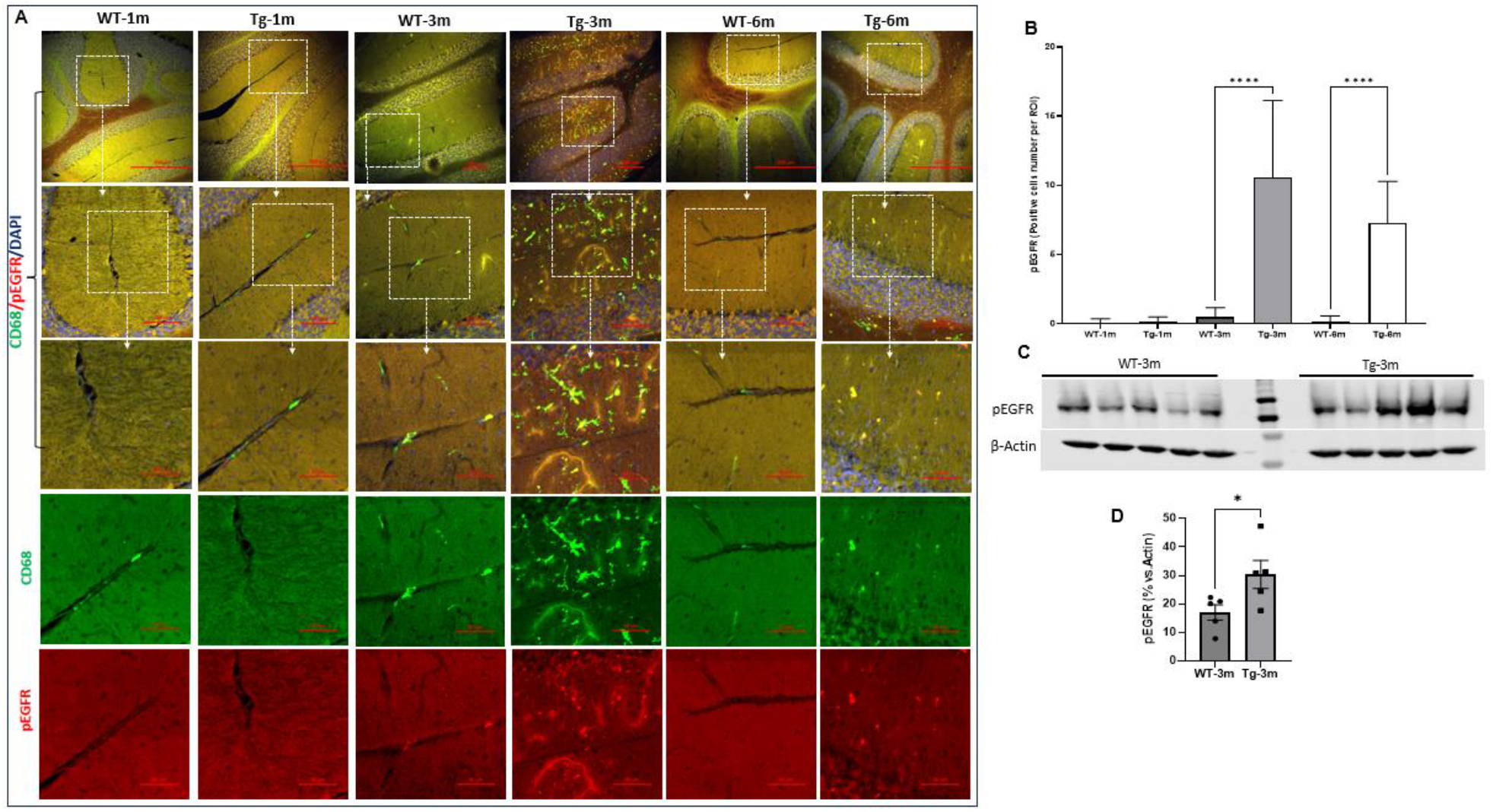
R424H mutation induced cerebellar pEGFR overexpression in macrophage. To assess the expression of phosphorylated (Tyr1068) EGFR (pEGFR) in macrophage, the location and protein levels of pEGFR and CD68 were examined in the cerebella of mice at different time points. Half of cerebella of mice were fixed for immunofluorescence staining, other half of the cerebella were fresh frozen for western blot analysis. (A) Images representatives of anti- pEGFR and anti-CD68 at the ages of 1,3, and 6 months. (B) Quantitative analysis of positive cells number of immunofluorescence staining signal of pEGFR (D) Western blot images of pEGFR. (E) Quantitative analysis of western blot of pEGFR. Note: Data was presented as mean ± SEM and unpaired t-test or one-way ANOVA Turkey multiple comparison test was used for quantitative analysis in GraphPad Prism 10. * p <0.05, **** p <0.0001.

### Correlation analysis indicated strong relationships between R424H mutation induced inflammation and neurodegeneration

To determine the relationship between inflammation and neurodegeneration in this R424H transgenic mouse model, we did correlation analysis between EGFR/pEGFR, CD68, GFAP, Iba-1 and Calbindin according to our quantitative expression data. We found there were strong inverse correlations between Calbindin and EGFR (r= -0.869, p=0.000), pEGFR (r= -0.817, p=0.004), and CD68 (r= -0.765, p=0.010) (Fig 10. A-C), but not GFAP (r= -0.268, p=0.455), or Iba-1 (r= -0.214, p=0.553) (Fig 10. D, E). This data suggested that increased CD68, EGFR, and pEGFR expression was associated with Purkinje cells loss in the cerebellum. Our correlation analysis also showed that there was a significant positive relationship between CD68 and EGFR/pEGFR (Fig 10. F, G), as well as between GFAP and pEGFR (Fig 10. H). This indicated that the expression of EGFR/ pEGFR was strongly related with inflammatory immune cells in the cerebellum.

**Fig 10.**
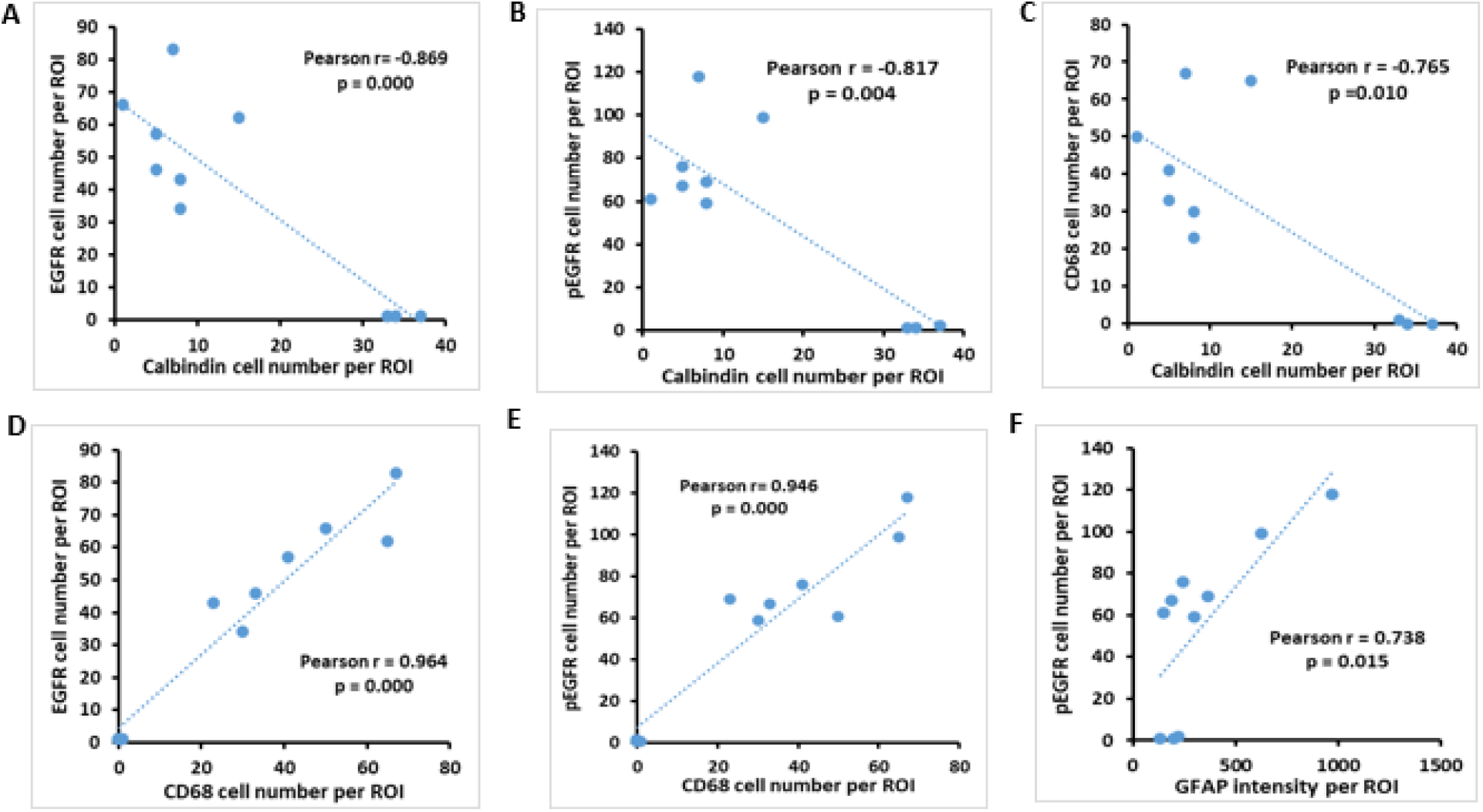
Correlation analysis indicated strong relationships between R424H mutation induced inflammation and neurodegeneration. Pearson correlation was analyzed among EGFR/pEGFR, CD68, GFAP, Iba-1 and Calbindin according to our quantitative expression data. (A) Correlation between EGFR positive cell number and Calbindin positive cell number. (B) Correlation between pEGFR positive cell number and Calbindin positive cell number. (C) Correlation between CD68 positive cell number and Calbindin positive cell number. (D) Correlation between EGFR positive cell number and CD68 positive cell number. (E) Correlation between pEGFR positive cell number and CD68 positive cell number. (F) Correlation between pEGFR positive cell number and the intensity of GFAP positive signal.

### R424H mutation affected peripheral inflammatory cytokines expression

Neuroinflammation is considered a common pathological characteristic of neurodegenerative disease given its role in aggravating neurodegeneration. Dynamic crosstalk between the peripheral immune system such as activated endothelial cells, monocytes, monocyte-derived dendritic cells, macrophages, T cells, and microglia and the central nervous system (CNS) has been considered as a major cause of neuroinflammation [59–61]. Our data showed there was a strong inflammatory response in the CNS of transgenic mice (Fig 5, Fig 6, Fig 7, Fig 8, and Fig 9). To better understand whether there is also an inflammatory response in the peripheral circulation system of transgenic mice, a battery of inflammatory cytokines was quantitatively measured using MSD-Plex Mouse Cytokine Kit. These cytokines are associated with inflammatory response and immune system regulation. These data indicated that there was no significant change observed between wild-type mice and R424H mice at the ages of 1 and 6 months. However, the plasma concentrations of IL-4, IL-5, IL-10, MCP-1, and MIP-2 in 3-month-old R424H mice were significantly lower in comparison to age-matched wild-type mice (p < 0.05 or p < 0.01), and the concentrations of IL-6 and IFNγ in 3- month-old R424H mice were significantly higher compared to wild-type mice (p < 0.05) (Fig 11). This data suggested that there was also a peripheral inflammatory response in these transgenic mice at 3 months of age.

**Fig 11.**
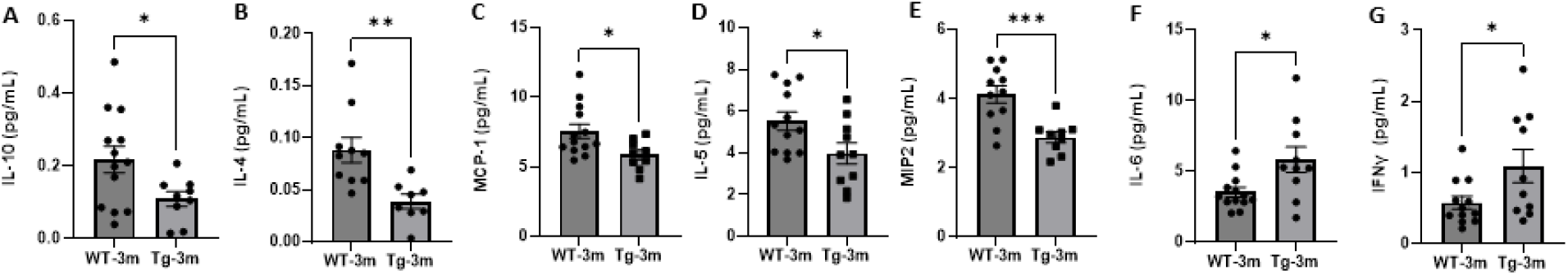
R424H mutation affected peripheral inflammatory cytokines expression. Plasma was collected from wild-type (WT) mice and R424H (Tg) mice at the ages of 1, 3, and 6 months. Plasma proinflammatory interleukins and cytokines were quantitatively measured using Meso Scale Discovery (MSD) V-PLEX Mouse Cytokine 19-Plex Kit according to the manufacturer’s protocol. (A) IL-10, (B) IL-4, (C) MCP-1, (D) IL-5, (E) MIP-2, (F) IL-6, and (G) IFNγ. Note: Data was presented as mean ± SEM and analyzed using unpaired t-test in GraphPad Prism 10. * p <0.05, ** p <0.01, *** p <0.001.

## Discussion

SCAs are a group of heterogeneous neurodegenerative diseases with phenotypes of cerebellar ataxia, extrapyramidal signs, dysarthria, oculomotor abnormalities, motor neuron signs, cognitive impairment, epilepsy, sensory deficits, and psychiatric manifestations [33, 62]. SCA13-R423H is a slowly progressive cerebellar ataxia with congenital onset mild cognitive impairment, dysarthria, nystagmus, and discrete pyramidal features [33, 63]. SCA13 mutations disrupt the firing properties of fast-spiking cerebellar neurons and influence neuronal function. Potassium channels (Kv3) are essential for fast spiking in burst neurons that fire hundreds of action potentials per second and Kv3 mutations cause abnormal neurodevelopment and neurodegeneration [33]. The double Kv3.1/Kv3.3 (encoded by KCNC1/KCNC3 genes) knockout mice showed marked symptoms, including tremor and severe ataxia, while single Kv3.3 knockout mice had no obvious motor phenotype [38, 64].

Here we have described a novel single SCA13 transgenic mouse model with a single *Kcnc3* R424H mutation. This model demonstrates motor neuron dysfunction with abnormal spontaneous firing of Purkinje cells and recapitulates the clinical phenotype of SCA13-R423H, in mirroring high-frequency tremor, aberrant gait, and the loss of coordination and balance.

Interestingly, the R424H mutation induced aberrant intracellular EGFR aggregation in our previous in vitro study [32]. In this study, we tested the expression of both EGFR and phosphorylated (Tyr1068) EGFR. Not surprisingly, both EGFR and pEGFR showed strong overexpression in the cerebellum, especially in the molecular layer. EGFR is known to be involved in growth, differentiation, maintenance, and repair of various tissues [45, 48], as well as a driver of tumorigenesis [46, 48]. Targeting overexpression of EGFR and associated signaling cascades are regarded as a rational and valuable approach in cancer therapy[46]. In adult and aged animals, the most prominent expression of EGFR is seen in cerebral cortex neurons [65]. For a long time, the role of EGFR in the nervous system was underestimated and poorly investigated [48]. However, recent studies showed that targeting EGFR expression had therapeutic benefits in neurodegenerative diseases and brain injuries [54–58]. The signaling activity of EGFR was related with its mobility and aggregation. EGFR aggregated in a phosphorylation-dependent manner [50]. Recent evidence also showed that aggregated EGFR further enhanced downstream phosphorylation and receptor endocytosis [51, 66] and exhibited neurotoxicity [51–53]. Our data showed that there was strong expression of EGFR/pEGFR in the molecular layer of the cerebella of 3 and 6-month-old R424H transgenic mice, which was minimally present in age-matched wild type mice. This data supports the hypothesis that over-activated EGFR/pEGFR signaling plays a critical role in R423H mediated cerebellar neurodegeneration. Further investigation is needed to determine whether EGFR/pEGFR signaling can be a potential therapeutic target for mitigating symptoms and pathology of SCA13 and beyond.

Recent studies reported that brain and spinal cord injuries upregulated EGFR/ pEGFR signaling and its downstream kinase activity in astrocytes resulting in the secretion of proinflammatory cytokines/mediators [54–58]. Cerebellar astrocytes were thought of as crucial players not only in the progression but also in the onset of distinct forms of ataxia [41]. In the cerebellum, astrocytes are classified into three main categories: Bergmann glia (BG) and granular layer astrocytes in the cerebellar cortex and fibrous astrocytes in the cerebellar white matter [67]. Hyperactive Bergmann glia could impair neuronal function by interfering with their normal supportive role or by triggering neuroinflammation [11]. Early astrocyte activation was indeed observed in both the cerebellar nuclei and cortex in a SCA21 mouse model [68]. A previous study in a SCA1 model showed that early activation of microglia and astrocytes were closely related with the onset and severity of SCA1[20]. The hyperactivated immune cells in the cerebellum exhibited obvious hypertrophy of cell bodies and processes, as well as increased expression of pro-inflammatory mediators in several SCA1 mouse models [20]. Interestingly, these activated cells were found well before the first signs of neurodegeneration and motor impairments in the affected cerebellar regions. This suggested that the mutation induced immune cell activation rather than activation initiated by neurodegeneration [20]. Importantly, the regulation of astrocyte reactivity mitigated neuroinflammation and degeneration of Purkinje cells, rescuing the motor deficits in mouse models of SCA1 and SCA6 [27, 69]. In our study, the data clearly demonstrated similarities in that there were hyperactivated astrocytes, microglia, and macrophages in the molecular layer of the cerebella in R424H transgenic mice. The inflammatory response reached its peak at 3 months in the transgenic mice. The activation of these immune cells and their EGFR/pEGFR signaling was strongly correlated with the loss of Purkinje cells. The damage triggered by these hyperactivated immune cells showed irreversible neurodegeneration in the cerebella of these transgenic mice. This suggested that early intervention for these immune cells’ activation may be a potential therapeutic strategy for these diseases. However, further experiments regarding the roles of certain types of immune cells in the neurodegenerative process are required.

Mounting evidence supports the existence of bidirectional crosstalk between the peripheral immune cells and the central nervous system [59–61, 70]. In this study, we found that the peripheral inflammatory cytokines such as IL-4, IL-5, IL-6, IL-10, MCP1, MIP-2, and INFγ, were also influenced in our transgenic mouse model. This significant change was only observed in 3-month-old transgenic mice. This change corresponded with the strongest observed inflammatory response within the CNS which also occurred at the 3-month age point. Blood brain barrier breakdown is usually hypothesized to be a common mechanism of dysregulated peripheral -CNS interplay, however the mechanism of crosstalk between the CNS and the periphery in this family of diseases requires further investigation. In conclusion, we have created a novel SCA13 transgenic mouse model which shows significant motor neurological dysfunction including high-frequency tremor, aberrant gait, and reduced vestibular function, thus mimicking the clinical phenotype of the disease in humans. This new model additionally exhibited obvious and progressive Purkinje cells loss and cerebellar atrophy. Importantly, this study demonstrated a strong inflammatory immune cell (astroglia, microglia and macrophages) activation in this mouse model. The activated immune cells potentially triggered EGFR/pEGFR overexpression and aggregation within the cells. This neuroinflammation was significantly related with neurodegeneration, Purkinje cells loss, and eventually cerebellar atrophy. Further investigation is required to validate the amplification of neuroinflammation by immune cells of CNS and periphery which may play an important role in the neurodegenerative mechanisms of SCAs.

## Abbreviations

CC3: Cleaved Caspase 3
CD68: Cluster of Differentiation 68 CNS: Central nervous system
EGFR: Epidermal growth factor receptor GFAP: Glial Fibrillary Acidic Protein GRO: human growth regulated oncogene
Iba-1: Ionized calcium binding adaptor molecule 1 IFN γ: Interferon gamma
IL-10: Interleukin-10 IL-12: Interleukin-12 IL-15: Interleukin-15 IL-17: Interleukin-17
IL-1β: Interleukin-1 beta IL-2: Interleukin-2
IL-27: Interleukin-27 IL-30: Interleukin-30 IL-33: Interleukin-33 IL-4: Interleukin-4 IL-5: Interleukin-5 IL-6: Interleukin-6 IL-9: Interleukin-9
IP-10: Interferon gamma-induced protein 10 KC: Keratinocyte Chemoattractant
KCNC3: Potassium Voltage-Gated Channel Subfamily C Member 3 MCP-1: Monocyte chemoattractant protein 1
MIP-1α: Macrophage inflammatory protein 1 alpha MIP-2: Macrophage inflammatory protein 2
MRI: Magnetic Resonance Imaging MSD: Meso Scale Discovery
NGS: normal goat serum
PBS: phosphate buffered saline pEGFR: phosphorylated EGFR R423H: pArg423His
R423H: pArg424His ROI: Region of interest
SCA: Spinocerebellar ataxias SCA13: Spinocerebellar ataxia 13 Tg: transgenic
TNF-α: Tumor necrosis factor alpha WT: wild type

## Declarations

**Ethics approval and consent to participate**: Not applicable.

**Consent for publication**: Not applicable.

**Availability of data and materials**: The datasets used and/or analysed during the current study are available from the corresponding author on reasonable request. The detailed sequence information during this study are included in this published article and its supplementary materials file.

**Competing interests**: The authors declare that they have no competing interests.

**Funding**: This study is funded by Seed fund at Mount Sinai (#02482323, MFW) and Barrow Neurological Foundation (#3033227, MFW). The generation and development of the CRISPR/Cas9 mutant mouse model was supported by the Jackson Laboratory National Institutes of Health Precision Genetics Center Grant (#U54OD020351, ARZ and CML).

## Acknowledgements

We acknowledge the support of Dr. Rick Maser and The Jackson Laboratory Genome Engineering Technology group. Core facility costs were defrayed by Cancer Center Support Grant (#CA034196) to The Jackson Laboratory.

## Authors’ information

Authors and Affiliations

Junxiang Yin, Michael Wu, Michael F. Waters.

Department of Neurology, Icahn School of Medicine at Mount Sinai, New York,10029, and Departments of Neurology, Barrow Neurological Institute, St. Joseph’s Hospital and Medical Center, Dignity Health, Phoenix, Arizona 85013, USA.

Jennifer White, Ming Gao

Departments of Neurology, Barrow Neurological Institute, St. Joseph’s Hospital and Medical Center, Dignity Health, Phoenix, Arizona 85013

Swati Khare,

Department of Neuroscience, University of Florida, Florida 32611, USA and Oxford PharmaGenesis Inc, Newtown, Pennsylvania 18940, USA

Jerelyn A. Nick, Harry S. Nick

Department of Neuroscience, University of Florida, Florida 32611, USA Aamir R. Zuberi5,

Technology Evaluation and Development, The Jackson Laboratory, Bar Harbor, Maine 04609, USA Cathleen M. Lutz

Rare Disease Translational Center, The Jackson Laboratory, Bar Harbor, Maine 04609, USA Kyle D. Allen

Department of Molecular Genetics and Microbiology, University of Florida, Florida 32611, USA.

## Authors’ contributions

JXY: Conceptualization, conceived, designed, performed, and analyzed data from all experiments, and wrote the paper. JW: review & editing, tissue harvest, mounted the sections to be observed under confocal microscope, acquired, and analyzed images. SK: review & editing, performed, and analyzed data of the MRI and motor function. MW: review & editing, mice tissue harvest, acquired, and analyzed images. ARZ: review & editing, contributed with animal model generation. MG: contributed with electrophysiological recorded and analyzed. JAN: performed and analyzed data of the MRI and motor function. CML: contributed with animal model generation, Funding acquisition. KDA and HSN: review & editing, supervised gait data. HSN: review & editing. MFW: Conceptualization, conceived, designed and supervised project, review & editing, Visualization, Funding acquisition. All authors discussed and approved the final manuscript.

## Corresponding authors

Correspondence to Junxiang Yin, email address, or Michael F. Waters, email address.

